# Identification of porcine RUNX1 as an LPS-dependent gene expression regulator in PBMCs by Super deepSAGE sequencing of multiple tissues

**DOI:** 10.1101/713206

**Authors:** Tinghua Huang, Min Yang, Kaihui Dong, Mingjiang Xu, Jinhui Liu, Zhi Chen, Shijia Zhu, Wang Chen, Jun Yin, Kai Jin, Yu Deng, Zhou Guan, Xiali Huang, Jun Yang, Rongxun Han, Min Yao

**Author notes:** Corresponding author’s contact information: College of Animal Science, Yangtze University, Jingzhou, Hubei 434025, China.

## Abstract

Genome-wide identification of gene expression regulators may facilitate our understanding of the transcriptome constructed by gene expression profiling experiment. These regulators may be selected as targets for genetic manipulations in farm animals. In this study, we developed a gene expression profile of 76,000+ unique transcripts for 224 porcine samples from 28 normal tissues collected from 32 animals using Super deepSAGE (serial analysis of gene expression by deep sequencing) technology. Excellent sequencing depth has been achieved for each multiplexed library, and principal component analysis showed that duplicated samples from the same tissues cluster together, demonstrating the high quality of the Super deepSAGE data. Comparison with previous research indicated that our results not only have excellent reproducibility but also have greatly extended the coverage of the sample types as well as the number of genes. Clustering analysis discovered ten groups of genes showing distinct expression patterns among those samples. Binding motif over representative analysis identified 41 regulators responsible for the regulation of these gene clusters. Finally, we demonstrate a potential application of this dataset to infectious and immune research by identifying an LPS-dependent transcription factor, runt-related transcription factor 1 (RUNX1), in peripheral blood mononuclear cells (PBMCs). The selected genes are specifically responsible for the transcription of toll-like receptor 2 (TLR2), lymphocyte-specific protein tyrosine kinase (LCK), vav1 oncogene (VAV1), and other 32 genes. These genes belong to the T and B cell signaling pathways, making them potential novel targets for the diagnostic and therapy of bacterial infections and other immune disorders.

## Introduction

The domestic pig (Sus scrofa) is an important farm animal for meat source worldwide and has been used as alternative models for studying genetics, nutrition, and disease as reviewed recently (Houpt et al. 1979; Verma et al. 2011; Bailey and Carlson 2019). The swine genome community has created a large amount of useful data about the transcriptome of pigs (Schroyen and Tuggle 2015). The recently released pig genome sequence (Sscrofa 10.2) (Groenen et al. 2012) and associated annotation greatly enhance our knowledge of the pig biology (Dawson et al. 2013; Beiki et al. 2019). Currently, it is estimated that the porcine genome encodes for ⍰ 40,000 genes (Groenen et al. 2012). Transcriptome analysis indicated that the actively transcribed genes are only a fraction, perhaps 15,000 genes, in normal tissues (Hornshoj et al. 2007). Several research groups have created microarray transcriptome profiling data for normal human tissues (Haverty et al. 2002; Shmueli et al. 2003), normal mouse tissues (Su et al. 2002; Su et al. 2004), and normal rat tissues (Walker et al. 2004). In the pig, several Expressed Sequence Tag (EST) sequencing projects, microarray platforms, and deep sequencing projects have developed gene expression profiles across a range of tissues (Hornshoj et al. 2007; Freeman et al. 2012). Compared to the model organisms, the information of the pig transcriptome is still limited in terms of comprehensive tissue and gene coverage (Schroyen and Tuggle 2015). Here we present Super deepSAGE (serial analysis of gene expression by deep sequencing) profiling data for the normal pig tissues with wide gene coverage and annotation. Using K-means clustering analysis and motif binding site enrichment analysis, we have identified regulators for co-expressed genes. A detailed analysis of one of the interesting transcription factors, runt-related transcription factor 1 (RUNX1), illustrated the power of the data.

## Results and discussion

### Development of the Super deepSAGE technology

A flowchart of the Super deepSAGE experiment is summarized in Fig. 1. Dynabeads® M-270 Amine (Thermo Fisher Scientific, China) were coupled with –C6-SH labeled reverse transcription-primer with the sequence containing the 5’-CAGCAG-3’ recognition site of EcoP15I and an Oligo(dT) sequence at 3’ end designed intentionally to complement the poly(A) sequence of mRNAs (Synthesized by Sangon Biotech, China). The coupling procedure was carried out following protocol reported by Hill and Mirkin (Hill and Mirkin 2006) using the succinimidyl 4-(p-maleimidophenyl)butyrate (SMPB) crosslink reagent (Thermo scientific, Shanghai, China). Ten micrograms of mRNA were reverse-transcribed (cDNA synthesis system, Invitrogen) with the Oligo(dT) magnetic beads to generate single-stranded cDNA using protocol recommended by the manufacturer. The product was converted to double-stranded cDNA using random primer and then digested with NlaIII (NEB, Beijing, China). The biotin-labeled linkers (linker-5EA) with phosphorylated 5’ termini and 3’ end overhang (5’-CATG-3’), containing the EcoP15I recognition site were prepared by annealing commercially synthesized oligonucleotides. The magnetic beads-bound cDNA was washed and linked to linker-5EA by T4 DNA ligase (NEB, Beijing, China). As a result, each cDNA fragment bounded to the magnetic beads is flanked by two inverted repeats of EcoP15I recognizing sites. The type III restriction enzyme EcoP15I cleaves the DNA downstream of the recognizing site (25 nt in one strand and 27 nt in the other strand) leaving a 5’ end overhang of two bases (Meisel et al. 1992; Moncke-Buchner et al. 2009). Linker-ligated cDNAs on the magnetic beads were digested with ten units of EcoP15I under conditions described previously (Matsumura et al. 2003). The supernatant containing released biotin-labeled fragments were added to streptavidin magnetic beads (Promega, Beijing, China), and the biotin-labeled fragments of the cDNAs were captured. Finally, barcoded linkers (linker-3EA) with two random base overhangs at 5’ end and phosphorylated termini were prepared and ligated to the cDNA ends by T4 DNA ligase (NEB, Beijing, China). The resulting products were amplified by polymerase chain reaction (PCR), and the 119 bp product was separated by polyacrylamide gel electrophoresis (PAGE) and recovered from the gel. The barcoded libraries prepared from different samples were combined into a single multiplex sequencing reaction at the end of library construction and submitted for deep sequencing. The sequence information of synthetic oligos, linkers, and primers are available in Supplemental document 1.

**Fig. 1.**
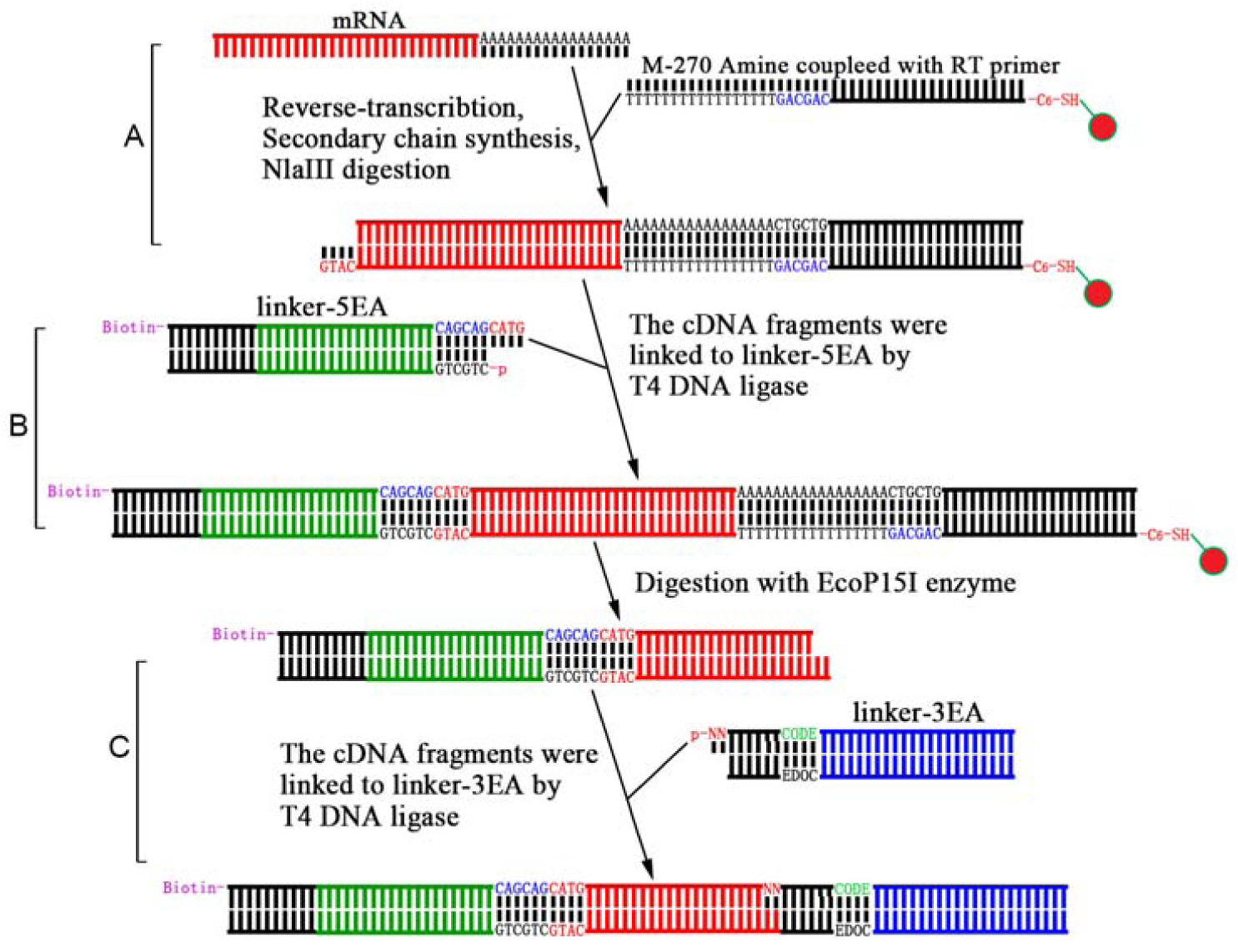
Flowchart of Super deepSAGE library construction. There are three major steps included in the protocol: A) reverse transcription with oligo(dT) coupled magnetic beads, synthesis of the secondary chain, and digestion with NlaIII; B) add 5’ end linker and digest with EcoP15I, and C) add 3’ end linker and PCR amplification. For details see materials and methods section.

The serial analysis of gene expression (SAGE) was first developed by Velculescu et al. (Velculescu et al. 1995) and improved by Saha et al. (Saha et al. 2002), Matsumura et al. (Matsumura et al. 2003), and Nielsen et al. (Nielsen et al. 2006). The traditional SAGE library construction protocol includes multiple steps, and the separation of the linker-tag fragment is challenging to perform, and the PAGE purification often produces low yield. The library construction protocol in this study was improved by introducing two magnet beads: 1) Dynabeads^®^ M-270 Amine coupled with –C6-SH labeled Oligo(dT) reverse transcription primer; 2) The streptavidin magnetic beads which can capture biotin-labeled linkers (linker-5EA). The magnetic beads used in this protocol can capture and purify the DNA fragments and is technically less demanding than PAGE separation. This modification increased the yield of linker-tag fragments and resulted in the robustness of the technique. Also, the primers and linkers were designed compatible with multiplexed deep sequencing technology, saving the sequencing cost.

### Animals, samples collection, and deep sequencing

A total of 224 tissue samples across 28 different tissues were collected from a slaughtering farm located in Hubei province in China. The samples were collected from 32 animals from a Duroc × Landrace × Yorkshire (DLY) commercial crossbreed pig populations consisting of 16 males and 16 females with a median age of 21 days. The endometrium, placenta, and conceptus were collected from Landrace × Yorkshire (LY) sows of 65 days of gestation. The detailed sample information is available in Table 1. In the computational extraction of tags from sequence data, the in-house designed program removes the two bases at the 5’ end. This ‘digital removal’ is performed to minimize the less accurate effect of two random bases, at the 5’ end of linker-3EA, and could potentially reduce the length of tags, and affect the representative ability of the data. However, direct link with a linker that has two random bases at the 5’ end forming stick ends will 1) enhance the efficiency of the link assay, and 2) no additional blunt ending process was needed. The inaccuracy caused by this linkage process was removed by the ‘digital removal’ procedure, thereby lowering the systematic bias in the data.

**Table 1.** Detailed information of the collected samples.

| ID | Code | Duplicates | Age (day) | Male/female | Breeds | Tissue |
| --- | --- | --- | --- | --- | --- | --- |
| 1 | AC | 8 | 21 | 4/4 | DLY | Adrenal cortex |
| 2 | AM | 8 | 21 | 4/4 | DLY | Adrenal medulla |
| 3 | CPT.SPH | 8 | 21 | 4/4 | DLY | Conceptus spherical |
| 4 | CPT.TUB | 8 | 21 | 4/4 | DLY | Conceptus tubular |
| 5 | FT.AB | 8 | 21 | 4/4 | DLY | Abdominal fat tissue |
| 6 | MS.DI | 8 | 21 | 4/4 | DLY | Diaphragm muscle |
| 7 | STOM | 8 | 21 | 4/4 | DLY | Stomach |
| 8 | CPT.FIL | 8 | 21 | 4/4 | DLY | Conceptus filamentous |
| 9 | MS.LD | 8 | 21 | 4/4 | DLY | Longissimus dorsi |
| 10 | LNG.TRA | 8 | 21 | 4/4 | DLY | Lung porcine trachea |
| 11 | PLACT | 8 | 21 | 4/4 | DLY | Placenta |
| 12 | LNG.BRO | 8 | 21 | 4/4 | DLY | Lung porcine bronchus |
| 13 | LNG.DIS | 8 | 21 | 4/4 | DLY | Lung porcine distal |
| 14 | SPL | 8 | 21 | 4/4 | DLY | Spleen |
| 15 | FT.BF | 8 | 21 | 4/4 | DLY | Back fat tissue |
| 16 | KID | 8 | 21 | 4/4 | DLY | Kidney |
| 17 | ADE | 8 | 21 | 4/4 | DLY | Adenohypophysis |
| 18 | MP.BMD | 8 | 21 | 4/4 | DLY | Bone-marrow derived<br>macrophage |
| 19 | MS.BF | 8 | 21 | 4/4 | DLY | Biceps femoris |
| 20 | EDMT | 8 | 21 | 4/4 | DLY | Endometrium |
| 21 | BLD | 8 | 21 | 4/4 | DLY | Blood |
| 22 | PBMC | 8 | 21 | 4/4 | DLY | Peripheral blood<br>mononuclear cell |
| 23 | HT | 8 | 21 | 4/4 | DLY | Heart |
| 24 | CC | 8 | 21 | 4/4 | DLY | Cerebral cortex |
| 25 | MP.MD | 8 | 21 | 4/4 | DLY | Monocyte derived<br>macrophage |
| 26 | MP.ALV | 8 | 21 | 4/4 | DLY | Porcine alveolar<br>macrophages |
| 27 | MLN | 8 | 21 | 4/4 | DLY | Mesenteric lymph nodes |
| 28 | LIV | 8 | 21 | 4/4 | DLY | Liver |

### Analysis of the complexity and diversity of Super deepSAGE data across tissues

Rarefaction analysis of size-fractionated library for each sample was performed to determine the complexity and diversity of the tissues in pig (Wang et al. 2018). The sequencing depth achieved using eight samples-multiplexed deep sequencing technic reached near-saturation of transcript discovery within all size ranges. Saturation was reached very early in Super deepSAGE sequencing data due to the lower complexity of the tags (number of tags) in libraries (Fig. 2A-F showed the first six deep sequencing runs). Samples from the same sequencing run were compared using reads from different size-fractionated libraries to further investigate the diversity of the relationship between sequencing depth and transcript discovery. In all deep sequencing runs, tissues exhibited transcriptome diversity in terms of both total numbers of reads and the number of transcripts discovered. For example, the muscle tissue (MS.DI_2) saturated much sooner than the conceptus (CPT.SPH_8) and have less number of transcripts discovered in the first deep sequencing run (Fig. 2A). Similar sequencing depth and diversity were obtained using size-fractionated data from each of the sequencing run and transcript as outcome measures (Supplemental Fig. SA-D).

**Fig. 2.**
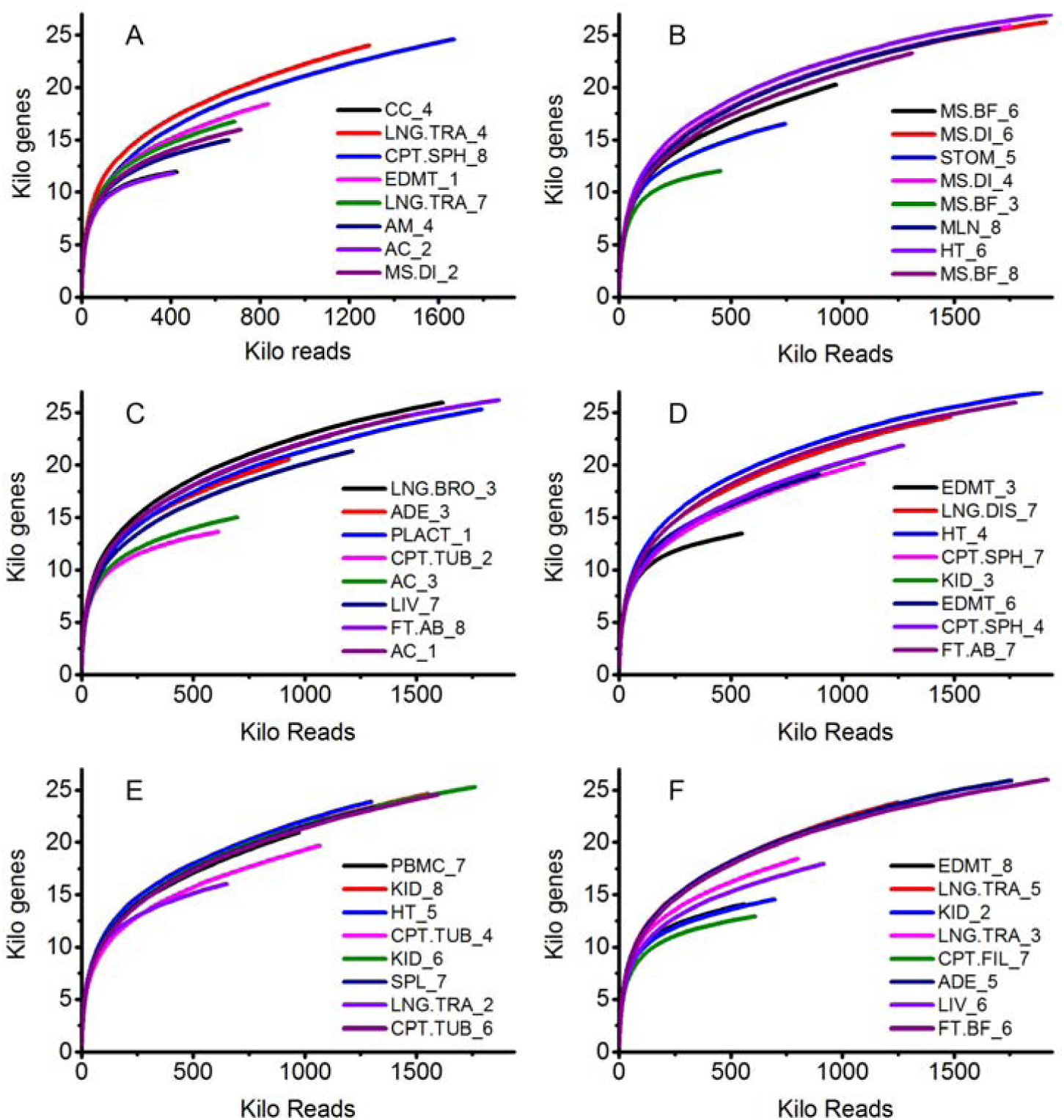
Rarefaction analysis of covered genes/transcripts in porcine tissues and cells Super deepSAGE library. Plot A to F shows the covered Kilo transcripts per Kilo reads in the first six Super deepSAGE sequencing runs. The samples in each sequencing run were randomized and detailed information is given in Table 1.

### Data quality and internal consistency control using principal component analysis (PCA)

Principal component analysis (PCA) was used to check if the samples clustered together according to their tissue source (Son et al. 2005). Even though the samples were collected from 32 individual animals from different families, genders, and ages (Table 1), the PCA plot showed that the samples from the same tissues clustered together and were distinct from other samples (Fig. 3). The transcripts in conceptus, blood, and macrophages had relatively distinct expression profile and segregation apart from the rest of the samples when plotted using the first two components of the PCA analysis (Fig. 3A). The adenohypophysis, cerebral cortex, heart, and muscle were aggregate and separated from other samples when plotted using the third and fourth component (Fig. 3B). The adrenal, liver, mesenteric lymph nodes, peripheral blood mononuclear cell, and spleen were slightly away from other samples when plotted using the fifth and sixth component (Fig. 3C). When removing those samples from the datasets and re-calculating the PCAs, the remaining samples; fat, placenta, endometrium, kidney, lung, and stomach grouped differently according to the tissue/cell types (Fig. 3D to F). Tissues having similar cellular composition and biological function, like alveolar macrophages and monocyte-derived macrophages or heart and skeletal muscles, clustered closely together but were separated from each other.

**Fig. 3.**
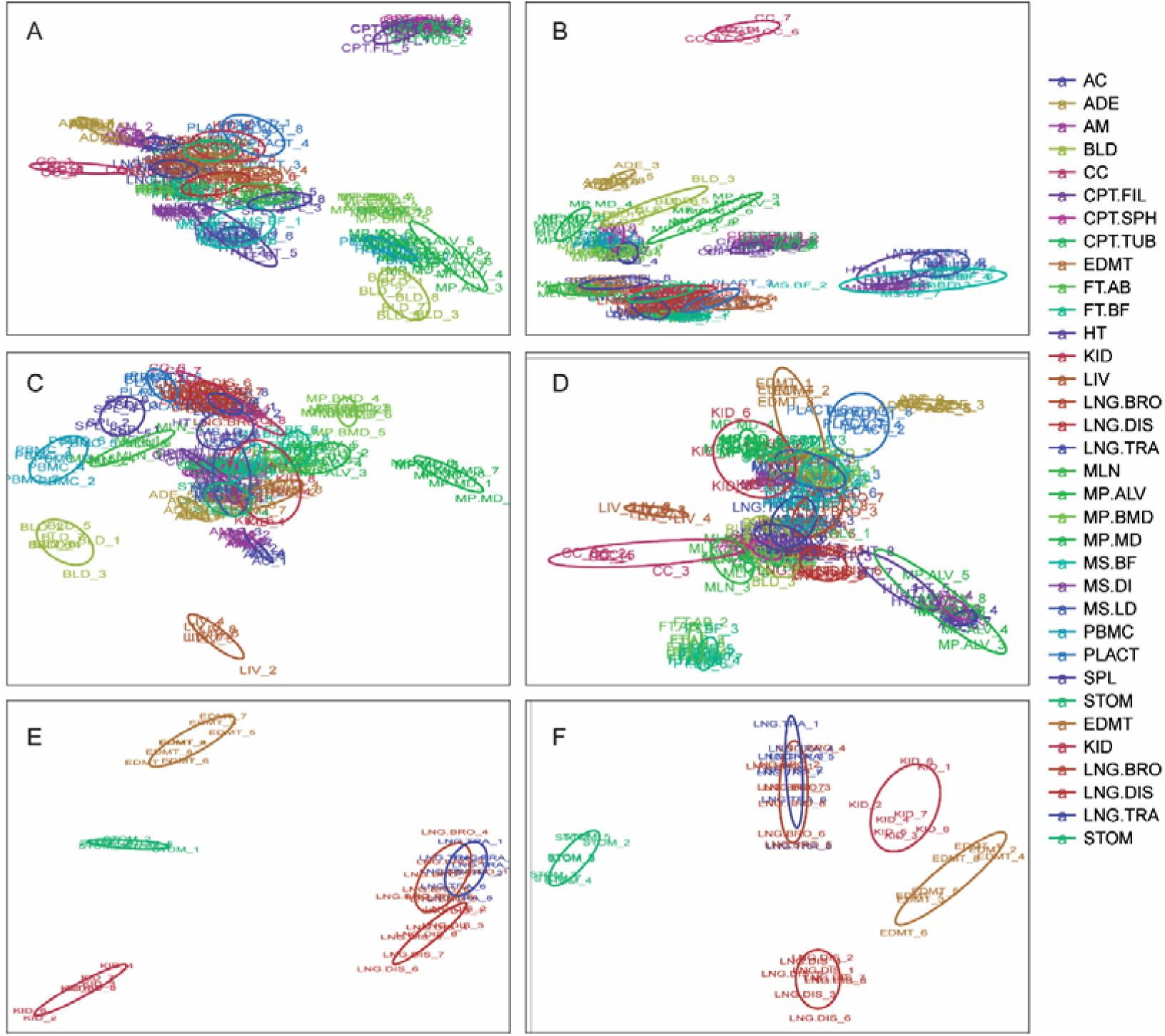
Principal components analysis of the Super deepSAGE sequencing data. A) to D) shows the top eight principal components of all 224 samples from the 28 tissues (two principal components per each plot). Samples separated in plot A to D were removed, and PCA was re-calculated with the remaining samples (fat, placenta, endometrium, kidney, lung, and stomach grouped). E) and F) shows the top four principal components of all the remaining samples (two principal components per each plot).

### Comparison of the Super deepSAGE data with previously published microarray research

The expression profiles were compared with microarray data published previously (Hornshoj et al. 2007). There is a total of 18,306 common genes for seven tissues, while high correlations (r=0.85-0.93 and p-values less than 1.0×e^-30^) were calculated between the gene expression profiles generated by the two platforms (Fig. 4). Similar dynamic range was observed in both platforms for transcripts with relative expression level between 0.55 and 0.95. Differences in expression profiles were apparent between the two platforms with several genes exhibiting relatively higher or lower expression values in either platform deviated from the diagonal line (Fig. 4). All transcripts had an expression value in the microarray, due to background hybridization or noise, regardless of whether it was truly expressed or not. The overall dynamics of the fitted curve showed that the Super deepSAGE is more sensitive than that microarray for the low expressed genes showing a concaved trend at the lower ends (with relative expression level less than 0.55 in Fig. 4). For those genes with high expression levels, variability is high in both Super deepSAGE and microarray platforms.

**Figure 4.**
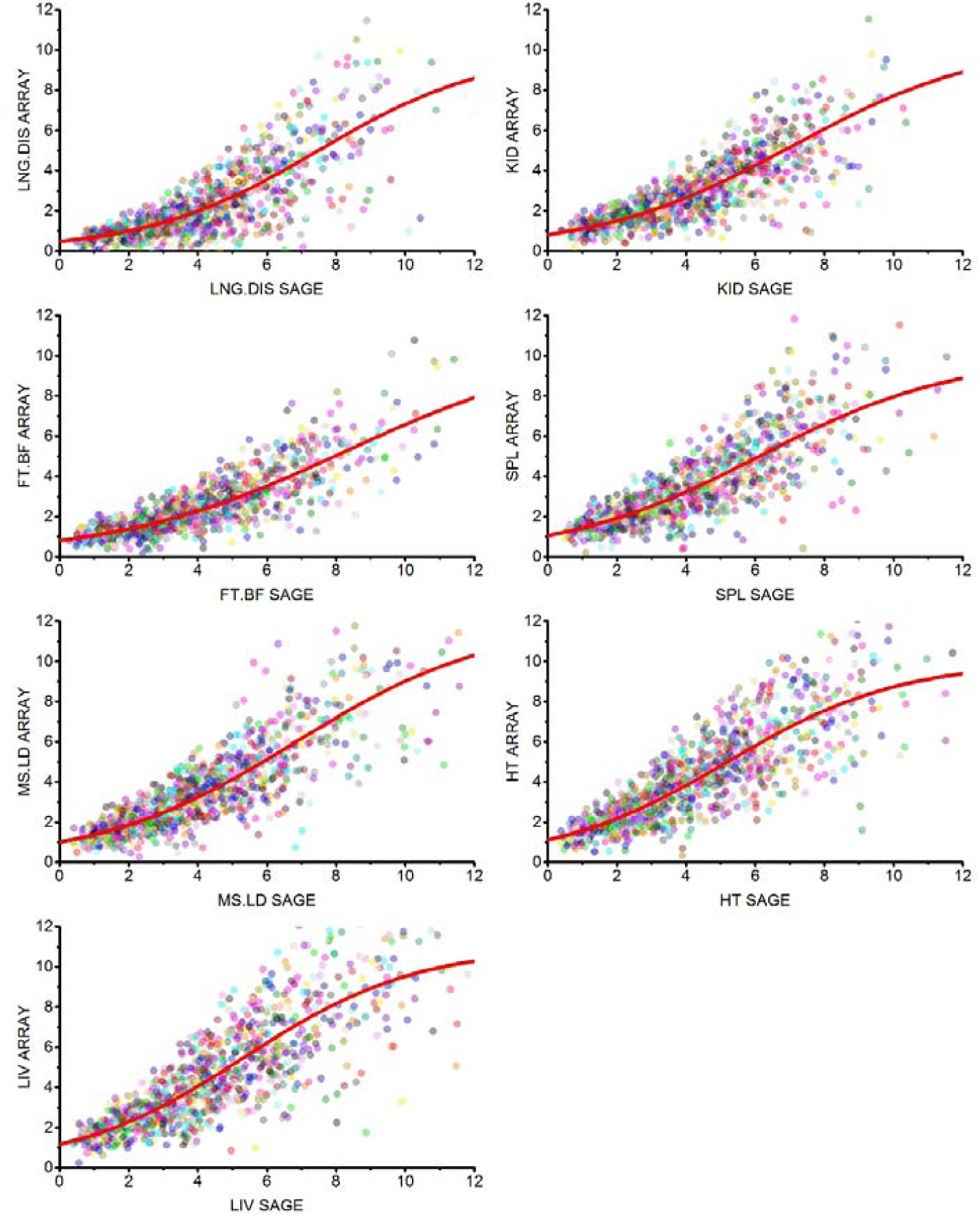
Comparison of the expression profiles of the 18,306 common transcripts between Super deepSAGE and microarray platforms. Scatter plots show the averages (between biological duplicates) of log_2_ transformed expression values of transcripts between two platforms. The relationship between the expression profiles generated in the two platforms is depicted as a smoothing spline (red).

As compared by microarray, reliable gene expression profiles can be generated by Super deepSAGE in seven known tissues. Of the 50 highest expressed Super deepSAGE tags, 38 (76%) found corresponding probesets in the 50 highest expressed genes, and only three tags showed a statistically significant difference between Super deepSAGE and microarray data. Two possibilities could cause such discrepancies between Super deepSAGE and microarray data: 1) the SAGE tag was derived from two or more different transcripts, which were differentially expressed in the samples tested, and 2) the microarray probeset can target two or more transcripts due to sequence similarity of transcripts. For example, the transcripts from the same gene family will always produce the same SAGE tag (attributable to the lower resolution power of Super deepSAGE) and preferred to hybrid to the same microarray probeset (can be minimized by design probesets in the none-conserved region). Regardless of some discrepancy, we conclude that Super deepSAGE data are overall compatible with the microarray data and provide faithful gene expression profiles.

### Identification of tissue-specific expression of transcripts

A total of 4,165 transcripts showed significant up or down-regulation at least in one tissues in comparison to the average tag count for all other 27 tissues. K-means clustering was then performed by trying a different number of centers (K from 5 to 28) and several random sets (S from 10 to 1000). Finally, we selected K = 10 and S = 400 to produce clustering result with clean and clear expression pattern (by visualization), highly reproducible for each duplicated run (Fig. 5). The result indicated that Cluster 1 has the largest number of transcripts, and most of these transcripts were expressed low in tissues, except macrophages, PBMCs, blood, and conceptus which were moderately expressed. The conceptus specifically expressed transcripts were in Cluster 2, while the conceptus, macrophages, PBMCs, and blood de-expressed transcripts were in Cluster 4. The macrophages, PBMCs, blood, mesenteric lymph nodes, and spleen specific expressed transcripts were in Cluster 5. The genes specifically expressed in heart and skeletal muscle were in cluster 10. The cerebral cortex specifically expressed genes were in Cluster 6, and liver specifically expressed transcripts were in Cluster 7. The adrenal cortex, adrenal medulla, cerebral cortex, and adenohypophysis specifically expressed transcripts were in Cluster 8. Transcript in Cluster 3 and Cluster 9 were ubiquitously expressed or expressed in multiple tissues.

**Fig. 5.**
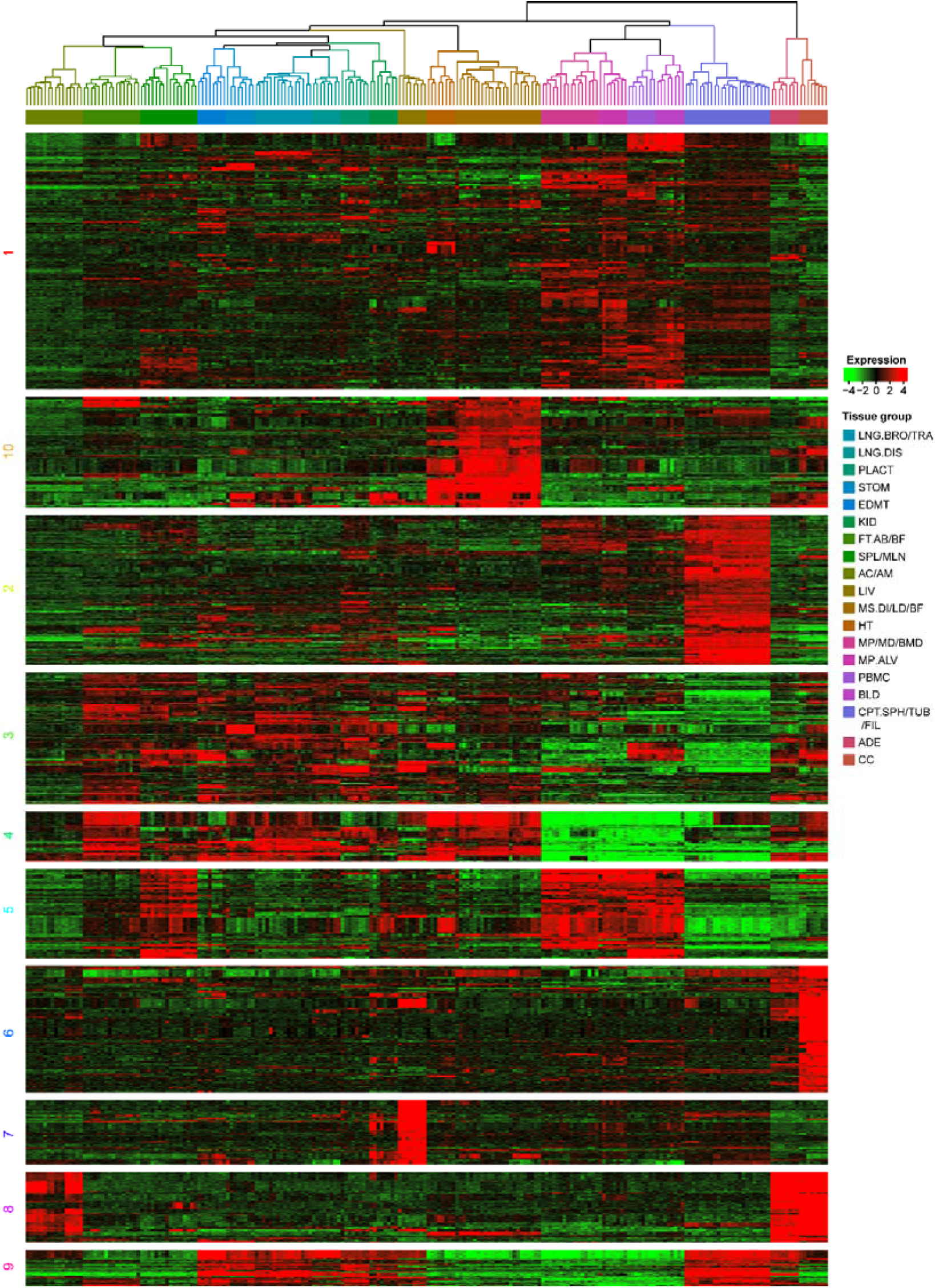
K-means clustering analysis of differentially expressed genes across tissues. Data adjustment (median center and normalization) was performed before the clustering analysis. The color codes of red, white, black, and dark green represent high, average, low, and absence of expression, respectively. A detailed view of expression pattern and internal structure of each gene cluster were constructed by hierarchical clustering and is shown in plot areas from 1–10.

Gene expression data obtained from transcriptional profiling experiments have inspired several applications, such as the identification of differentially expressed genes (Huang et al. 2011; Huang et al. 2018) and the creation of gene classifiers for improved diagnoses of diseases such as cancer (Wesolowski and Ramaswamy 2011; Tonella et al. 2017). The gene expression profile of 224 samples created in this study is complicated that traditional models were difficult to apply to this data to find differentially expressed genes. An *ad hoc* method comparing each tissues to the average tag count for all other 27 tissues was performed, and a very stringent threshold was set (fold change >5.0, p-value <1.0×10^-6^) to filter the tissues specifically expressed transcripts. The K-means clustering algorithms which group similar transcripts and separate dissimilar transcripts by assigning them to different clusters have proven to be useful for identifying biologically relevant gene clusters for different biological status (Yao et al. 2016). Even though very useful, the K-means clustering algorithm is particularly sensitive to initial starting conditions and converges to the point that is the local minimum (Selim and Ismail 1984). Furthermore, the number of clusters (parameter K) is difficult to be determined. In this study, global-seeding procedures of BF98 (Bradley and Fayyad 1998) have been introduced into the algorithm to improve the consistency and quantity of clustering results. The BF98 method employed a bootstrap-type procedure to determine the initial seeds for the centers. Several subsamples (recommended n = 10) of the data set were clustered using K-means. Each clustering operation produced a different candidate set of centroids from which a new data set was constructed. This data set was clustered using K-means, and the centroids were chosen as the initial seeds. The optimal BF98 clustering result on the Super deepSAGE data was obtained by “visualization” of the result performed by using K=10 and number of subsamples S=1000 after trying K from 5 to 28 and S from 20 to 1,000. The “visualization” method is straightforward for that deterring the best parameter for the K-means clustering procedure, but when the K reached 10, definite, compact and representative gene clustering was formulated, and when the S is higher than 200, consistent clustering result was produced for each duplicated clustering run.

### Identification of over-represented motif for tissues specifically expressed transcripts

The CLOVER software (Frith et al. 2004) with JASPAR PWM database (Khan et al. 2018) was used to identify over-represented transcription factor binding motifs for each cluster of genes. The promoter regions for each cluster of transcript (1,000 bp upstream) were obtained using the Ensemble Biomart tool (Kasprzyk 2011). The promoter regions for the whole transcript detected in this project, which possesses similar GC content, were used as background. Motifs having a p-value of ≤ 0.05 were selected as significant (Table 2, top 5 motifs). The most significantly enriched motif in Class 1 is MZF1. TFAP2A and TFAP2C were also significantly enriched with a raw score higher than 30. In Class 2, there was only one significantly enriched motif, RHOXF1. In Class 3 and 4, there were five and four motifs with p-value < 0.05 respectively, but the raw score was lower than ten. In Class 5, there were at least five motifs with p-value < 0.05, and three of them, RUNX1, ASCL1, and Myod1 had a raw score higher than 30. In Class 6, the significantly enriched motifs with the highest score were SNAI2 and FIGLA, whereas, in Class 7, the significantly enriched motifs with the highest score was NR4A2. In Class 8, there was only one motif ZEB1 enriched in the promoter region of these transcripts. In Class 9, all the enriched motifs had a raw score of less than ten. In Class 10, the top three motifs were Ascl2, Myog, and Tcf12.

**Table 2.**
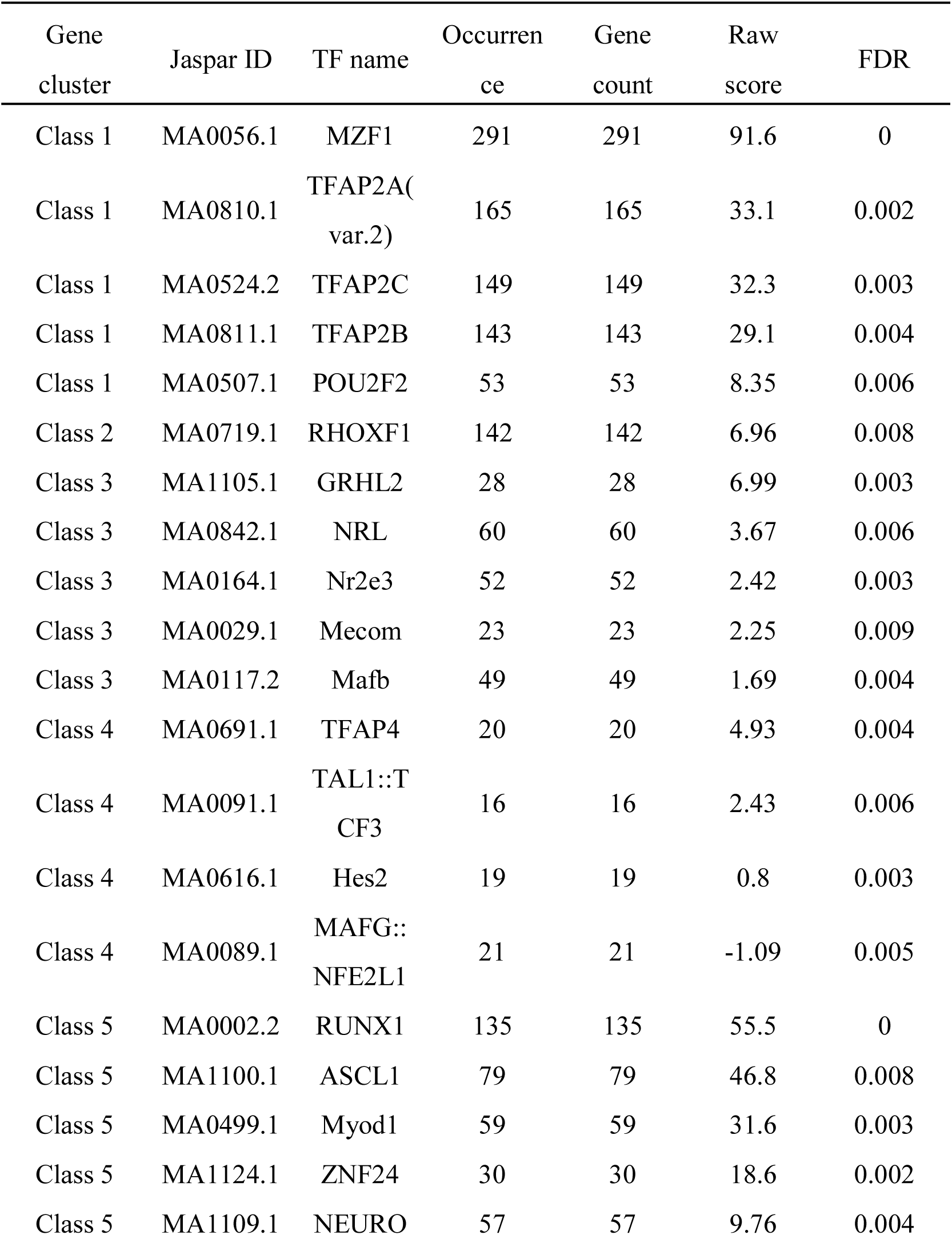
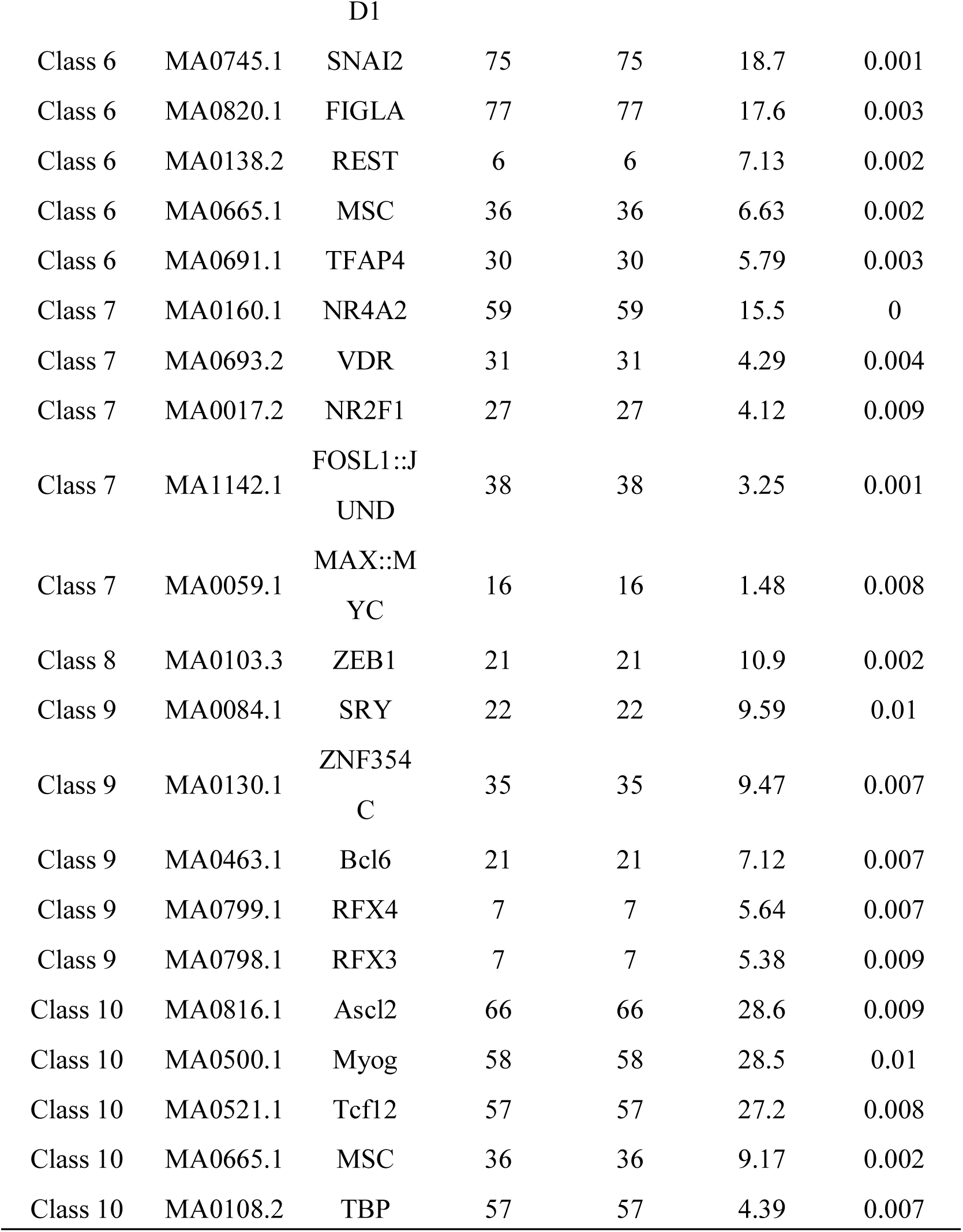
Significantly over-represented binding motifs in the promoter region of transcripts showing a similar expression pattern

The transcription factors interact with the DNA recognition motifs, regulates transcription of a large number of genes, and play important roles in fundamental biological processes, including growth, development, and disease (Latchman 1997). To understand gene expression regulation in the Super deepSAGE data obtained in this study, identifying the over-represented or under-represented motifs in the sequence showing similar expression pattern and which factors bind to them, is necessary. Over-representation indicated the motif candidates playing a regulatory role in the sequences, while under-representation indicated that the motif would have a harmful dis-regulatory effect. In each gene clusters showing a similar expression pattern, Clover successfully detects motifs known to function in the sequences and generate interesting and testable hypotheses.

### Case report: confirmation of the regulatory roles of RUNX1 in PBMCs in pig

#### Confirmation of the RUNX1 binding site in the promoter region of TLR-2, LCK, and VAV1

The toll-like receptor 2 (TLR-2), lymphocyte-specific protein tyrosine kinase (LCK), and vav1 oncogene (VAV1) plasmid containing the 1Kb promoter sequence were used in *in vivo* studies (wild type). To show the regulation effect of RUNX1, the binding site of RUNX1 in TLR-2, LCK, and VAV1 was mutated or deleted. Reporter vectors constructed by the wild type, mutated, or deleted promoter sequences were transfected into the peripheral blood mononuclear cells (PBMCs), and luciferase activity was monitored. Binding site deletion significantly attenuated the expression of the downstream reporter luciferase activity (p<0.05), indicating that the RUNX1 could interact with the target site and regulate the expression of the downstream reporter gene (Fig. 6A-C). The mutated vectors showed significant attenuation of the activity of downstream luciferase at 40, 44, and 48 hours post-transfection (p<0.05) indicating a regulatory relationship between RUNX1 and the targets. Another experiment was performed using mouse macrophage cells (RAW 264.7) to validate the hypothesis further. Consistent with the previous results, deletion/mutations to the RUNX1 binding sites in TLR-2, LCK, and VAV1 promoter sequence significantly attenuated the activity of downstream luciferase at 40, 44, and 48 hours post-transfection (Fig. 6D-F). The luciferase reporter activity after transfection with the wild-type vector was significantly higher in macrophage cells than in the PBMC assays, suggesting that the endogenous RUNX1expression in mouse macrophage cells was higher than in PBMCs.

**Fig. 6.**
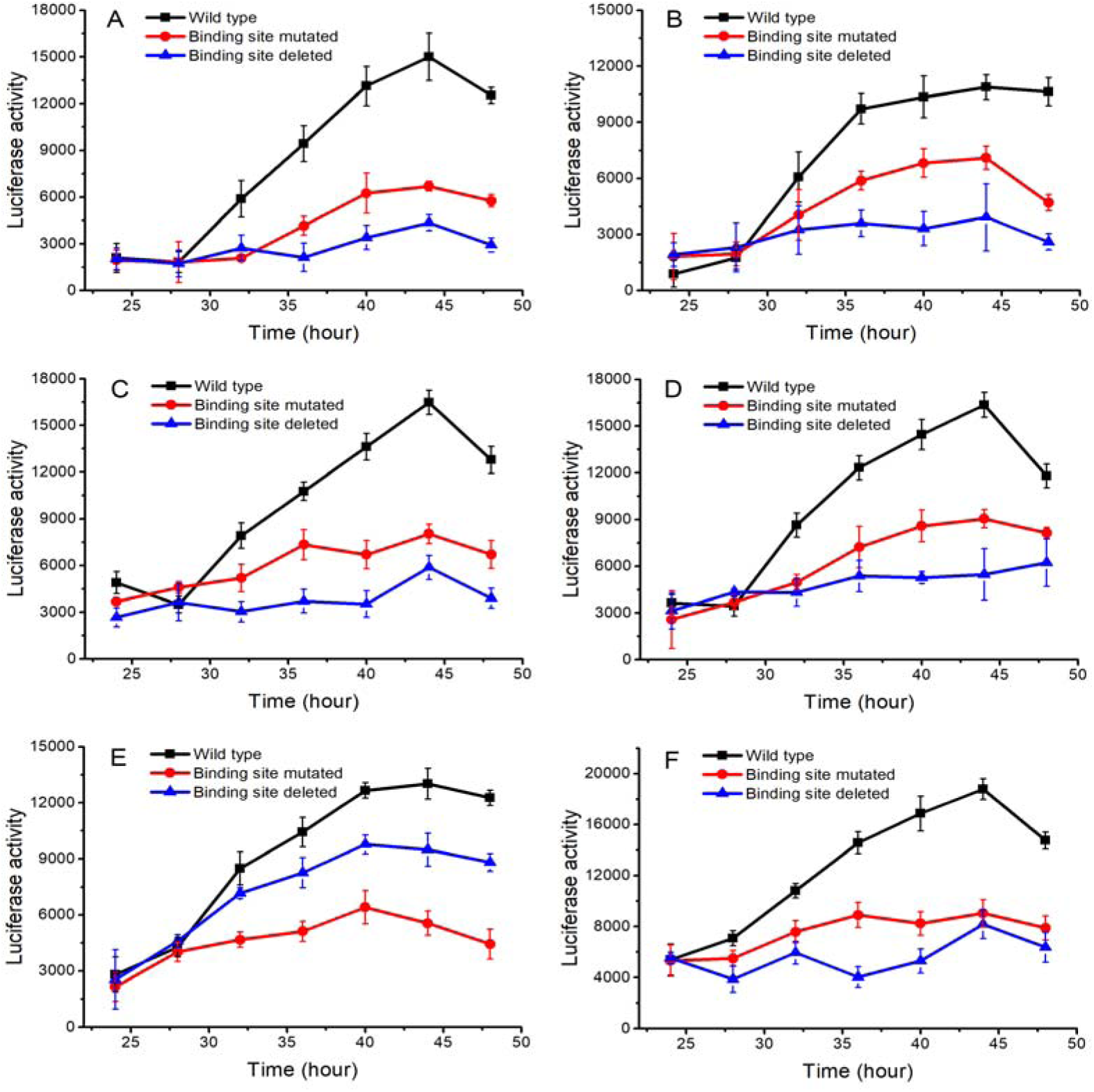
Luciferase reporter assay of the RUNX1 targets. One wild-type promoter construct (containing the predicted RUNX1 binding site), two mutant constructs (mutated or not containing the binding site) were investigated. The mutant construct (black) was identical to the wild-type, except that the RUNX1 binding site was deleted or mutated. The line graphs show the luciferase activity after the reporter plasmids were transfected into PBMCs (A-C) or macrophages (D-F). Three RUNX1 target gene have been investigated (A and D: TLR-2, B and E: LCK, C and F: VAV1). The error bars represent the mean ± standard deviation of three duplicate samples.

### RNA flow cytometry analysis of RUNX1 targets in LPS and RUNX1 inhibitor treated PBMCs

To show the effect of RUNX1 on three targets; TLR2, LCK, and VAV1, pig PBMCs were stimulated with LPS and/or RUNX1 inhibitor, for 6 hours, during which their TLR2, LCK, VAV1, CD14 protein levels were monitored. Two subsets of cells readily emerged from CD14/TLR2 analysis in PBMCs: a CD14^hi^/TLR2^lo^ (CD14^high^/TLR2^low^) and a CD14^lo^/TLR2^lo^ population (Fig. 7D). The percentage of CD14^hi^/TLR2^lo^ cells increased in LPS plus RUNX1 inhibitor treated samples, but the proportion of CD14^lo^/TLR2^lo^ cells remained unchanged. The percentages of TLR2^hi^ (for both CD14^hi^ and CD14^lo^) cells increased seven-fold in LPS alone treated samples compared with the non-treated controls. Four subsets of cells readily emerged from CD14/LCK analysis in PBMCs treated with LPS or RUNX1 inhibitor: a CD14^hi^/LCK^lo^, CD14^hi^/LCK^hi^, CD14^lo^/LCK^hi^, and CD14^lo^/LCK^lo^ population (Fig. 7E). The percentage of CD14^hi^/LCK^hi^, and CD14^lo^/LCK^hi^ cells increased in LPS plus RUNX1 inhibitor treated samples, and the proportion of CD14^hi^/LCK^lo^ cells was decreased. The percentages of CD14^hi^/LCK^hi^ cells increased by 40% in LPS alone treated samples compared with the non-treated controls. Two subsets of cells readily emerged from CD14/VAV1 analysis in PBMCs: a CD14^hi^/VAV1^lo^ and a CD14^lo^/VAV1^lo^ population (Fig. 7F). The percentage of VAV1^hi^ (for both CD14^hi^ and CD14^lo^) cells increased four-fold in LPS plus RUNX1 inhibitor treated samples. The percentages of VAV1^hi^ (for both CD14^hi^ and CD14^lo^) cells increased seven-fold in LPS alone treated samples compared with the non-treated controls and is two-fold higher than in LPS plus RNUX1 inhibitor treated samples.

**Fig. 7.**
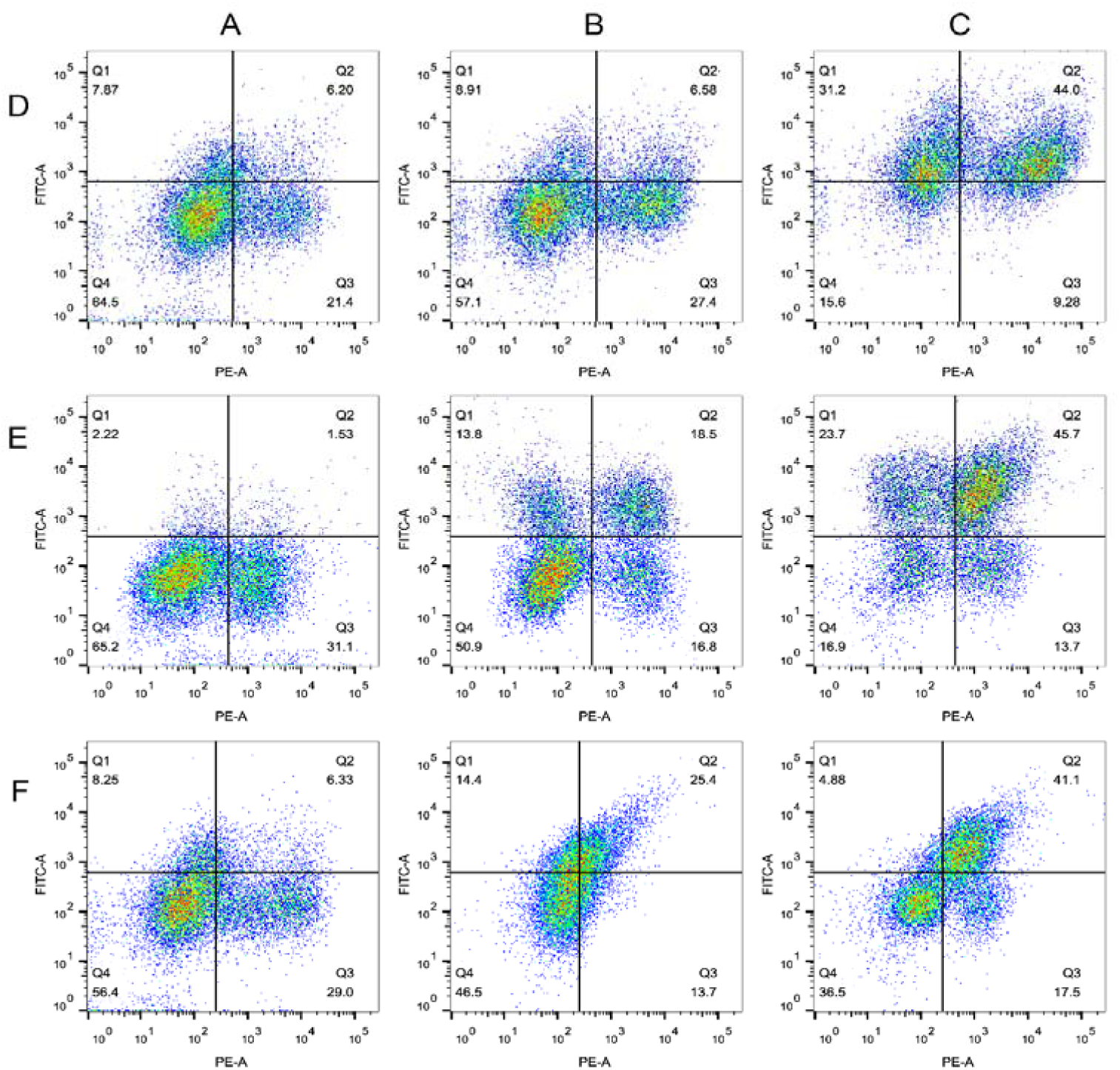
Simultaneous staining of the target gene and CD14 protein in rested and stimulated PBMCs. Plots of PBMCs that were left untreated (A) or were stimulated with LPS plus RUNX1 inhibitor for 6 hours (B) or were stimulated with LPS only (C), and labeled with antibodies that bind to CD14 (PE-A) and target protein (FITC-A, D-F).

RUNX1 is a master regulator of hematopoiesis and plays a vital role in T and B cells development. RUNX1 is critical in inducing the production of genes in immune cells, such as interleukin-2 (IL-2, (Wong et al. 2011), IL-3 (Uchida et al. 1997), colony-stimulating factor 1 receptor (CSF1R, (Zhang et al. 1996), CSF2 (Frank et al. 1995), and cluster of differentiation 4 (CD4, (Taniuchi et al. 2002). However, its roles in LPS-mediated inflammation in PBMCs remains unclear. In this study, regulations of TLR-2, LCK, and VAV1 have been confirmed by flow cytometry. TLR2 is an essential receptor for the recognition of a variety of pathogen-associated molecular pattern (PAMPs) from Gram-positive bacteria, including bacterial lipoproteins, lipomannan, and lipoteichoic acids (Medzhitov 2001). LCK encoded protein is a key signaling molecule in the selection and maturation of developing T-cells (Davis and van der Merwe 2011). The VAV1 encoded protein is important in hematopoiesis, playing a role in both T-cell and B-cell development and activation (DeFranco 2001; Helou et al. 2015). These results suggested that RUNX1 might be a new potential target for resolving inadequate or uncontrolled inflammation in PBMCs.

### Real-time PCR analysis of RUNX1 targets in LPS and RUNX1 inhibitor treated PBMCs

To investigate if the expression patterns of the 23 RUNX1 target genes could be modeled by LPS and RUNX1 inhibitor treatment *in vivo*, we performed real-time PCR assay after treating PBMCs with two different doses of LPS (1 ng/mL, 10 ng/mL), and RUNX1 inhibitor (1 ng/mL, 10 ng/mL). Samples were collected six hours post-stimulation. A total of 21 genes were induced in response to at least one dose of LPS stimulation, as expression levels for these genes were different when compared to non-stimulated control. A total of 10 genes were down-regulated in response to the RUNX1 inhibitor treatment. Hierarchical clustering analysis was used to determine whether the response of LPS stimulation response was similar to the patterns detected in RUNX1 inhibitor treatment, and if any differences were observed depending on the dosage of LPS and RUNX1 inhibitor used. As shown in Fig. 8, the expression patterns of samples with RUNX1 inhibitor treatment, RUNX1 inhibitor plus LPS treatment, and non-simulated controls clustered together. Different dose of the RUNX1 inhibitor did not affect the samples, as observed by the mixing up of respective samples in the heatmap. The LPS treated samples were unique and were separated from the RUNX1 inhibitor-treated groups and control groups. Similar to the RUNX1 inhibitor, different doses of LPS dose did not affect the samples as well. The expression patterns of RUNX1 inhibitor plus LPS treatment samples were similar with controls and samples treated with RUNX1 alone because they were mixed in the heatmap.

**Fig. 8.**
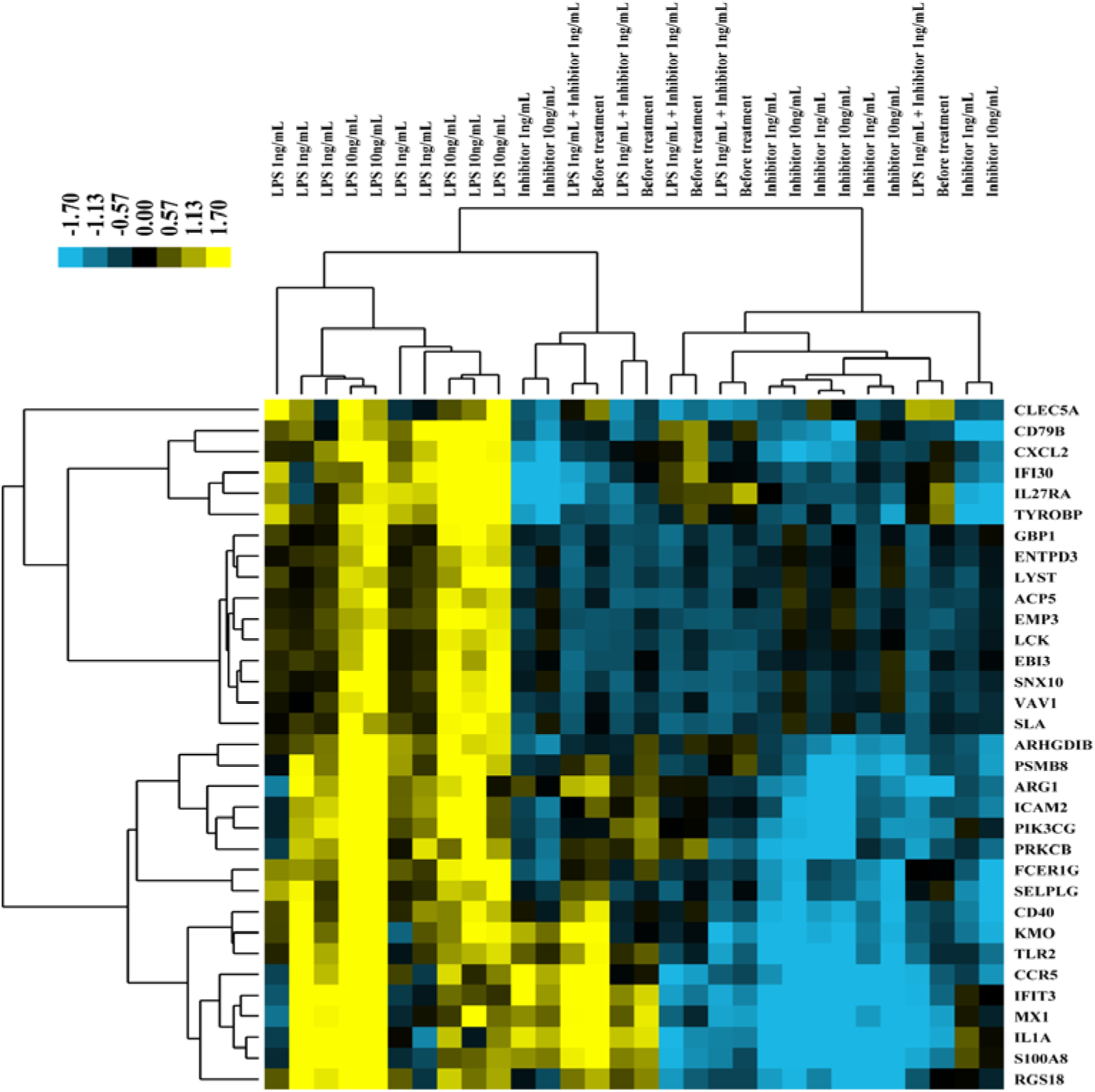
RUNX1 target gene expression in PBMCs treated with LPS or RUNX1 inhibitor. Cells were treated in vitro with two different doses of LPS (1 ng/ml, 10 ng/ml) and RUNX1 inhibitor (1 ng/ml, 10 ng/ml). Color codes of yellow, black, and blue represent expression levels of high, average, and low, respectively, across the treatments shown.

### Super deepSAGE is a useful data resource in pig study

Gene expression analysis is extensively applied in the understanding of the molecular mechanisms underlying a wide range of biological process such as host-pathogen interactions. Our dataset of transcript levels in normal tissues was developed as a reference datasets that can be compared to attained information of biological event specifically related aberrations in transcript levels. Therefore, one major focus of this manuscript was to demonstrate the biological importance of these profiles. We report that >40% of the measured transcripts were differentially expressed between the different tissues. We show that statically the transcripts were co-regulated by a few important transcription factors. We describe one of the many transcription factors that regulated gene expression in PBMCs. To our knowledge, this data set is the largest to date for the analysis of transcriptional profiles within normal tissues from pigs and is complementary to previously published data sets. These data will improve the annotation of the pig genome, support versatile biological research, and increase the utility of the pig as a meat source animal and model in medical research.

## Materials and methods

### Sample collection and RNA extraction

Soon after anesthesia by electric shock, specimens were excised, snap-frozen in liquid nitrogen, and kept in a deep freezer (−80°C) until RNA extraction. RNA extraction from the tissue samples and cells was conducted using the RNeasy Mini Kit (Qiagen, Shanghai, China) following the manufacturer’s protocols. The BioAnalyzer 2100 (Agilent) was used to assess the integrity of total RNAs, and RIN number of less than 0.7 was removed from the study.

### Super deepSAGE sequencing and data procession

The sequencing data were filtered by removing sequences that had poor quality (score <0.5) for more than 20% of all the bases. All the data discussed in this study have been deposited to the NCBI GEO database (Edgar et al. 2002) under accession number GSE134461. Tag sequence was extracted, counted, and assigned for each transcript, and then normalized for each sample by quantile normalization method (Pan and Zhang 2018). For the tags assigned to multiple transcripts, the average copy numbers of those tags were used. The principal component analysis (PCA) was performed using the log_2_ tag counts of all the transcripts across all the samples using R statistical software version 3.5. The tissue specific transcripts expressed were identified by comparing samples from each tissue to the overall tag count across all samples (average), and a threshold was set to fold change >5.0, p-value <1.0×10^-6^ according to a method implemented in limma package (Ritchie et al. 2015). Clustering analysis was performed by first using K-means clustering method to separate the transcripts to several big groups, and then using Hierarchical clustering to build the internal structure of the transcripts within the groups according to the method reported by Gu et al. (Gu et al. 2016).

### Luciferase reporter assay

The three predicted target genes, TLR-2, LCK, and VAV1, were also conserved in human and mouse. For these three genes, a 1Kb nucleotide promoter segment that included RUNX1 target sites was inserted upstream of a firefly luciferase ORF (pGL3, Promega, Beijing, China), and luciferase activity was compared to that of an analogous reporter with point substitutions disrupting the target sites, or analogous reporter that the binding site deleted completed (see detailed sequence information in Supplemental document 2). The logic behind the luciferase reporter assay is that deletion/mutation of a RUNX1 binding site should allow the down-regulation of its target genes, and hence the target gene should be expressed differently between the wild type and mutated constructs. The pGL3-Control activity was used for the normalization of firefly luciferase activity. For the assay, the cells were plated in a 96-well plate at 3,000 cells per well. After overnight incubation, the cells were treated with a transfection mixture consisting of 35 μL of serum-free medium, 0.3 μL of TransFast™ Transfection Reagent (Cat. E2431), and 0.02 μg of pGL3 and pGL3-Control vector per well. After one hour incubation, 100 μL of the serum-containing medium was added to the wells. At 24 to 48 hours of post-transfection, EnduRen™ Live Cell Substrate (Cat. E6481) was added to a final concentration of 60 μM, and luciferase activity was monitored.

### PBMC Isolation and Stimulation

Peripheral blood mononuclear cells (PBMCs) were isolated from whole normal blood collected from five animals aged 21 days using BD Vacutainer^VR^ Cell Preparation Tubes (Becton Dickinson, Shanghai, China). The samples were processed according to the manufacturer’s instructions within two hours of blood collection. PBMCs were harvested from the tube, washed with phosphate-buffered saline (Life Technologies), and centrifuged for 10 min at 300g prior to use. To induce gene expression, PBMCs were resuspended in RPMI-1640 medium (Life Technologies) supplemented with 10% fetal bovine serum (Life Technologies) at 1.5×10^6^ cells/mL in a 96-well V-bottom polypropylene plate (Corning Incorporated). LPS (Sigma-Aldrich, Shanghai, China) and RUNX1 inhibitor (Ro 5-3335, R&D Systems, Shanghai, China) were added at 5 ng/mL and 10 ng/mL, respectively, according to the manufacturer’s instructions. Untreated PBMCs were used as control samples.

### Surface staining and cytometry acquisition

Phenotypic surface staining was performed in BD Pharmingen™ stain buffer (BSA, BD Biosciences, Shanghai, China) for 30 min at room temperature in the dark, using anti-CD14 PE (BD Biosciences, Shanghai, China). Cells were washed and suspended in BD Pharmingen stain buffer (BSA, BD Biosciences, Shanghai, China), anti-TLR-2 FITC, anti-LCK FITC, anti-VAV1 FITC (BD Biosciences, Shanghai, China), was then added separately, and the mixture was incubated for 20 min at room temperature. Finally, cells were washed and acquired on a BD LSRFortessa™ cell analyzer (BD Biosciences, Shanghai, China). The flow cytometry data were deposited in Flow Repository database (ref) under accession FR-FCM-Z268.

### Data access

https://www.ncbi.nlm.nih.gov/geo/query/acc.cgi?acc=GSE134461

http://flowrepository.org/id/RvFrphtLijqf34kFNTA1gdB6BdXEskSDTdhZ4VwfM1qbgTIPfmqbL8o5eVTIhiUH

## Acknowledgments

This project was funded by the National Natural Science Foundation of China (NSFC Grant No. 31402055), the College Students’ Innovation and Entrepreneurship Training Program of Yangtze University (Grant No. 2018057), the Science and Technology Research Project of Department of Education of Hubei Province (Grant No. Q20171305), the Yangtze Youth Talents Fund (Grant No. 2015cqr12), the Yangtze Youth Fund (Grant No. 2015cqn39).

## Ethics approval and consent to participate

All procedures involving animals were ethical and were approved by the Animal Care and Use Committee of Hubei Province (China, YZU-2018-0031).

## Disclosure declaration

The authors declare that the research was conducted in the absence of any commercial or financial relationships that could be construed as a potential conflict of interest.

